# Sex specific correction of maternal inflammation-induced behavioral abnormalities by the inhibition of colony-stimulating factor 1 receptor

**DOI:** 10.1101/2025.05.30.657085

**Authors:** Jean Christophe Delpech, Hana Yeh, Srinidhi Venkatesan Kalavai, Yang You, Zhi Ruan, Nina Touch, Samuel Hersh, Pauline Monguillon, W Evan Johnson, Andrea Rau, Charlotte Madore, Tsuneya Ikezu, Seiko Ikezu

**Affiliations:** Department of Pharmacology and Experimental Therapeutics, Boston University School of Medicine, Boston, MA 02118, USA; Graduate Program in Neuroscience, Boston University, Boston, MA 02118, USA; University of Bordeaux, INRAE, Bordeaux INP, NutriNeuro, UMR 1286, F-33000 Bordeaux, France; Université Paris-Saclay, INRAE, AgroParisTech, GABI, 78350, Jouy-en-Josas, France; Division of Infectious Diseases, Center for Data Science, Rutgers University, New Jersey Medical School, Newark, NJ, 07103; Department of Neuroscience, Mayo Clinic, Jacksonville, FL 32224, USA

**Keywords:** behavior, colony stimulating factor 1 receptor, maternal immune activation, microglia, neurodevelopmental disorders, Polyinosinic:polycytidylic acid, regeneration, transcriptomics

## Abstract

We have previously reported the therapeutic effect of depletion and regeneration of endogenous microglia on autism spectrum disorder-like behaviors in offspring from the dams under maternal immune activation (MIA) by dampening their neuritogenic activation. Here we show a long-lasting pathological effect by MIA, leading to abnormal behaviors in offspring mice in a sex-specific manner at 3 months of age. MIA was induced by injecting Polyinosinic:polycytidylic acid [Poly(I:C)] at 10mg/kg at E9.5 with preselected dams based on the immunoreactivity to low-dose Poly(I:C) injection. MIA offspring show impairments in sociability, repetitive behavior, and spatial working and associative memories in both sexes, whereas only male offspring show social novelty deficit. MIA has no effect on anxiety-like behavior nor sensori-motor or locomotor activity. Administering colony stimulating factor receptor (CSF1R) inhibitor in young adult MIA offspring regenerates microglia and ameliorates sociability deficits in males and spatial working and associative memory impairments in both sexes without effect on social novelty impairments. Transcriptomic analysis of prefrontal cortex and hippocampal tissues reveals MIA and sex-specific pathways and highlights potential key genes involved in the corrective effect of microglial regeneration in MIA offspring. Our results underscore the potential of the sex-specific therapeutic applications of CSF1R inhibitor for MIA-related social and cognitive disorders.

## Introduction

Viral or bacterial infection during the first trimester of pregnancy has been associated with a higher incidence of neurodevelopmental disorders (1,2). Animal models of maternal immune activation (MIA) have been widely studied to understand their underlying pathogenesis. MIA is commonly induced in pregnant rodents by the viral mimetic polyinosinic:polycytidylic acid [Poly(I:C)] (3). We and others demonstrated that MIA offspring exhibit long-lasting behavioral abnormalities such as increased repetitive behavior as well as impaired social interaction and communication (3,4). In regards to MIA models, attention in the field has recently started to shift toward understanding resilience and susceptibility in large part due to the recurring issues of reproducibility in MIA models (5–9). As a result, the McAllister group demonstrated that maternal factors, even before pregnancy, strongly influence how exposure of isogenic mice to the same gestational risk factor leads to divergent outcomes in offspring (10). We thus tested the immune reactivity of the females before pregnancy, in order to enhance the reproducibility of MIA phenotypes in the offspring, while describing its differences from other MIA conditions. In addition, recent evidence, including ours, has shown the possible involvement of different immune cell types, including Th17 lymphocytes (11) and microglia (4), the primary innate immune cells in the central nervous system, as contributors of the MIA phenotype in offspring. In addition to acting as first immune responders in the brain, microglia also support synaptic maturation and circuit formation during critical developmental periods (12–15). We recently showed that colony stimulating factor-1 receptor (CSF1R) inhibitor-induced microglia renewal at adulthood was able to restore the MIA phenotype (4), highlighting the active role played by the microglia throughout adulthood. We demonstrated that CSF1R inhibitor treatment corrected microglial phenotypic changes and aberrant interactions with neurons, accompanied by normalized synaptic properties, spine morphologies and a complete reversal of MIA-induced abnormal social behavior in male offspring (4). Neurodevelopmental disorders are more frequently diagnosed in males than females over the course of late adolescence or early adulthood (16). Recently, the MIA model was used to study sex differences in the schizophrenia model, and a similar deficit of social behavior was found in both sexes, while only males showed sensorimotor gating deficits and anxiety-like behaviors (17). However, cognitive functions were not explored, even though neurodevelopmental disorders-associated behaviors (and/or other phenotypes?) were observed. Building on our recent data showing a promising effect of microglia renewal on the MIA phenotype in male offspring, we decided to test whether our CSF1R inhibitor treatment approach is able to correct the MIA phenotype in females by testing cognitive abilities in both sexes of MIA offspring and the potential beneficial effect of CSF1R inhibitor treatment on cognition under MIA condition.

Here we report that our MIA model induces sex-specific transcriptomic changes in the prefrontal cortex (PFC) and hippocampus (HPC). At the behavioral level, our optimized MIA model induces social, working memory and spatial associative memory abnormalities in both sexes, without affecting locomotor and sensory-gating abilities. In addition, we report that microglia depletion induced by administration of a CSF1R inhibitor followed by microglial repopulation reversed MIA-induced abnormal social behavior in males alone and reversed associative memory abnormalities in both sexes. Overall, CSF1R inhibitor treatment seems to be globally efficient in regards to memory abilities but male specific in regards to social abilities, in agreement with the recent line of research dedicated to delivering personalized medicine instead of a generalized treatment approach.

## Methods

### Animals

Male and female C57BL6 mice (C57BL6/N from Charles River laboratory and Taconic laboratories; C57BL6/J from Jackson laboratory) were kept on a 12-h light / 12-h dark schedule with access to chow *ad libitum*. Pregnant mothers were single-housed after one-night timed mating. Offspring were weaned at P21 and group-housed with 2-5 same sex littermates per cage. Male and female mice were used and litters for the MIA and saline groups were separated due to prenatal injection of pregnant mothers. The number, sex and age of animals for each experiment are listed in Figure legends. No statistical methods were used to predetermine sample size, and randomization of samples was not required. All animal procedures followed the guidelines of the National Institutes of Health Guide for the Care and Use of Laboratory Animals and were approved by the Boston University Institutional Animal Care and Use Committee.

### Maternal Immune Activation and Microglia Depletion/Repopulation

#### Blood collection and plasma preparation

For pre-selection and 3-hour timepoint post Poly(I:C) injection at E9.5, 50-200ul of blood was collected by venipuncture of the facial vein as previously described using 5 mm lancet (Goldenrod Animal Lancer, Medipoint, Mineola, NY (PMC4311747)). The dripping blood was collected into K3 a EDTA Sample Tube (MV-E-500, SAI Infusion Technologies), mixed by inversion of the tube, and placed on ice for transport. For IL- 17 at 48h after Poly(I:C) injection at E11.5 in Charles River and Taconic C57BL6 mice, blood was collected immediately after CO2 euthanasia by cardiac puncture with 1ml syringe precoated with EDTA (BD, Thermo Fisher). Blood was centrifuged at 2000 rcf for 10 minutes at 4C to collect blood plasma and stored at -80C until ELISA testing.

#### Maternal Immune Activation

Poly(I:C) Invivogen Tlr-pic (lot # PIC-40-07) was resuspended at concentration of 1 mg/ml in sterile saline. For preselection, eight-week-old females were intraperitoneally (ip) injected with HMW Poly(I:C) based on body weight (5mg/kg). Following at least one week of rest, nine- to twelve-week-old adult female C57BL6 mice were subjected to one night timed-mating with male C57BL6 mice (Charles River). Females were separated the following day, and checked for weight gain on E9.5 to determine pregnancy. On embryonic day 9.5 (E9.5), female mice were intraperitoneally (ip) injected with 10 or 20 mg/kg Poly(I:C) (Sigma, P0913, lot 128M4040V) as previously described (10). E9.5 corresponds to the middle of the first trimester in terms of human developmental biology, at which time studies have reported an increased likelihood of prenatal infection being associated with neurodevelopmental alterations in humans (1).

Lost pregnancy was confirmed by the presence of embryos in the abdomen after no birth/continuous weight gain. This paradigm has been confirmed to lead to physiological markers of immune response in mothers and microglial alterations in offspring (18).

#### Microglia depletion and Repopulation

Microglia depletion and subsequent repopulation was achieved through administration of the colony stimulating factor 1 receptor (CSF1R) inhibitors PLX5622 (1200 p.p.m. delivering daily doses of 168 mg/kg) (Plexxikon, Inc., mixed into AIN-76A standard chow by Research Diets, Inc.) from postnatal day P21-42 (MG-REP group) (19,20).

#### ELISA

Blood plasma harvested from mice was measured for plasma level of IL-6 and IL-17 1α. The cytokines IL-6 and IL17 were measured by the commercial enzyme-linked immunosorbent assay (ELISA) kits following the manufacturer’s instructions (Mouse IL-6 ELISA MAX™ Deluxe, 431304, Biolegend and Mouse IL-17 ELISA MAX™, 432504, Biolegend). Due to small collection volume from pre-selection and E9.5, we diluted blood plasma using 1x Assay Buffer provided by the manufacturer at a 1:20 ratio.Female responders were assigned to categories based on IL-6 quartiles (below the lowerquartile=low; between the second and third quartiles= middle; greater than the upper quartile= high). For IL-17, no dilution was used.

#### Behavioral Testing

All behavioral tests were performed in an empty testing room within the animal vivarium, during the light cycle between the hours of 9am-6pm. For all tests, animals were habituated in the test room for at least 1 hour before the start of testing. A test-free period of 5-7 d was used between behavioral tests. The lighting, humidity, temperature, and ventilation were kept as constant as possible in the testing room. The experimenter was not present in the room during any of the tests. The experimenter performing behavior tests was blind to the treatment group. All behavioral analyses were performed in a blinded manner. Animal sample sizes are reported in the Figure legends.

#### Three-Chamber Social Interaction Test

The social interaction test was performed as previously described (21,22). An opaque plexiglass three-chambered apparatus was custom-designed by our lab and fabricated by Boston University Scientific Instrument Facility. A mouse was placed in the middle chamber with door-blocking walls in place, and left to habituate for 5 min. Then, an unfamiliar same age, sex, and weight-matched mouse (within 5g) was placed in a clear perforated plexiglass cylinder in one of the side chambers, to the right or left of the entranceway. An identical empty cylinder was placed in the opposite side chamber. The chamber wall dividers were removed, and the test mouse was allowed to explore freely for 10 min while being video recorded, allowing sociability of the test mouse to be measured. After 10 min, a second mouse, identified as novel compared to the now familiar mouse present in one cylinder, was introduced in the empty cylinder and the test mouse was allowed to explore freely for an additional 10 min while being video recorded, allowing social memory of the test mouse to be measured. The mouse was then placed in a new empty cage and the apparatus was cleaned thoroughly with 20% ethanol. The time spent in each chamber and the number of entries into each chamber were video-tracked using Ethovision XT14 software.

#### Open Field test

Anxiety-like behavior and locomotor activity were assessed in a square open-field (40x40 cm) made of white coated plywood with 16-cm high walls. The floor was divided into center and periphery zones. Mice were placed in the central area of the apparatus and allowed to explore freely for 10 min. The total amount of time spent in the center is an index of anxiety-like behavior, while the total distance and the velocity are an index of locomotor activity.

#### Pre-Pulse Inhibition (PPI)

For PPI measurements, an SR-LAB system acoustic startle box with digitized electronic output (San Diego Instruments) containing a piezoelectric accelerometer mounted under a Plexiglas cylinder was used to generate and measure startle response and PPI. Each testing session began with a 5-min acclimation period at background noise intensity of 65 db. This was followed by four pulse alone trials, then a pseudorandom admixture of pulse-alone, pre-pulse + pulse, and no stimulus trials; followed by four more pulse alone trials for a total of 48 trials per test session according to a previously established protocol (23). For all trials, the SR-LAB machine was programmed to deliver acoustic startle stimuli (or no stimulus) over a background noise level of 65 db with a variable inter-trial interval; the startling stimulus was presented as a fast-rise noise burst lasting 40 ms at an intensity of 120 db. The animal’s whole-body flinch response to each stimulus was recorded as 48 consecutive 250-ms recordings beginning at stimulus onset. For pre-pulse + pulse trials, a pre-pulse of 3, 6, and 12 db over background (68, 71, and 77 db) of 20 ms preceded the primary pulse by 100 ms. Baseline startle reactivity was calculated from the average startle magnitude for the initial four pulse alone (120 db) trials from the first PPI session. Pre-pulse inhibition was defined as the percentage of the decline of startle response (pre-pulse inhibition (%) = 100−[(startle amplitude after pre-pulse and pulse)/(startle amplitude after pulse only)×100]).

#### Y-maze test

The Y-maze assesses working memory based on the innate preference of a mouse to alternate arms while exploring a new environment. Typically, mice prefer to explore a new arm of the maze rather than returning back to the one that was previously explored. The Y-maze apparatus consisted of three arms with dimensions 35cm (length) × 7cm (height) x 5cm (width) (San Diego Instruments). Testing was always performed in the same room and at the same time to ensure environmental consistency as previously described (24). In the training session, one arm was closed with a gate and a test mouse was placed at the end of one arm and allowed to explore the two opened arms freely through the maze during a 5-min session. Then, the mouse was removed, the closing gate removed and the maze cleaned using 20% ethanol. After a 30 min time inter-interval, the test mouse was placed back in the Y-maze and allowed to explore the entire maze freely during a 5-min test session. The total amount of time spent in each arm (starting-familiar-new) was tracked using Ethovision XT14 software, and time in familiar vs novel arm was used to calculate an index of recognition. The maze was cleaned with 20% ethanol after each mouse to minimize odor cues.

#### Novel object recognition

During the training phase, mice were introduced with two identical objects two times during 5 minutes, separated by a 5 minute interval (two training sessions). Animals exploring less than 2 sec with objects during the second 5 minute training session were excluded from the analysis. 24 hours later, during the testing day, mice were introduced first during 5 minutes with the same two familiar objects. After a 5 minute inter-trial interval, one object was replaced with a new object in the same arena at the same location as the familiar object, and exploration of the novel object was recorded during 5 minutes. Time spent by the mouse exploring each object during all the training and test sessions was measured manually by a blinded experimenter. Total time exploring the familiar and novel object during the final test session and an index of recognition calculated using the time exploring the familiar and the novel object during the final test session were reported.

#### Fear conditioning

The fear conditioning tests were performed as previously described (25,26) with minor modifications. Briefly, mice were trained and tested in a conditioning chamber (26 × 34 × 29 cm, Med-Associates Inc.) equipped with black methacrylate walls, a transparent front door, a speaker and grid floor. On day one, each mouse was placed into the conditioning chamber for a learning session. Baseline freezing was quantified during the initial 4-min period. Beginning at 4 min and at 120s intervals thereafter, the mouse was exposed 3 times to a 20 sec sound and a 2 sec 0.75-mA continuous foot shock overlapping the end of the sound (unconditioned stimulus; US). Broadband white noise was used instead of a frequency-specific tone in an effort to avoid possible auditory deficits that might occur with age. The mouse was removed from the chamber 1 min after the last foot shock, and placed back in its home cage. The contextual fear-conditioning memory was tested 24 h after the training phase, when the animal was placed back inside the conditioning chamber for 5 min without any shock. The freezing responses to the environmental context were quantified over a 5-min period to evaluate contextual fear conditioning. The cued fear-conditioning memory was tested 48 h after the training phase, when the animal was placed into a novel environment in the conditioning chamber and allowed to explore for 240 sec, at which point the same tone as day 1 was played for 60 sec, followed by 180 sec of no-tone (post-tone period). This was repeated two more times for a total time in the novel context of 960 sec.

### Immunohistochemistry

For cortical abnormalities during development, E18.5 embryonic mouse brains were post-perfused with 4% PFA. For adult microglia analysis, mice were transcardiacly perfused with ice-cold PBS followed by 4% PFA. Brains were then collected for post-fixation and cryoprotection with 30% sucrose/PBS. Brain sections were cut coronally on a cryostat and antigen retrieval with sodium citrate. Immunohistochemistry was performed to stain SATB2 (Abcam, ab51502), TBR1 (Abcam, ab31940), IBA1 (Wako, 019-19741). Sections were blocked in 5% BSA, 5% normal goat serum (NGS), and 0.2% Triton X-100 for 2 h. Sections were then incubated in primary antibodies (mouse anti-SATB2 1:1000; rabbit anti-TBR1 1:1000; rabbit anti-mouse IBA1 1:1000), in 1% BSA and 0.02% tween buffer over night at 4°C. Sections were then washed and incubated in secondary antibody (AlexaFlruor488 goat anti-mouse secondary antibody, 1:5000; AlexaFluor546 goat anti-rabbit secondary antibody, 1:5000, A- 11010) in PBS 1%BSA and 0.02% tween buffer for 2 hours. They were stained with Dapi for 2 min, washed 10 min in PBS, and mounted on slides with Prolong Gold antifade solution (Life Technologies, P36930).

### Imaging

Cortical abnormality images were captured by Nikon Eclipse epifluorescence microscope at 10x. Confocal imaging was performed on a Leica TCS SP8 lightning microscope at the inverted Leica DMi8 microscope stand using the confocal mode with HC PL APO CS2 20x/1.3 objective. Images of 2048 × 2048 pixels as confocal stacks with a z-interval of 0.20 μm system optimized was used to image tissue sections. For imaging Iba-1, a 488-nm laser line was used and emission was collected at 490-600 nm. Gain and off-set were set at values which prevented saturated and empty pixels. For microglial counting, Imaris software 9.5, 64-bit version (Bitplane AG, Saint Paul, MN) was used. Final data analysis was performed using Microsoft Excel and Graph rendering was done in GraphPad Prism.

### Image analysis

Automated detection of IBA1+ microglia density was performed using IMARIS (Bitplane). Images were tiled using SP8 Leica. Regions of interest (ROS) consisting of mPFC and hippocampus were manually selected for each brain section. The total density of microglia was calculated by dividing the microglia number by the ROI volume. A total of five to six regions were averaged per animal to acquire microglia density per volume.

### RNA sequencing

Brains were collected after cold PBS perfusion, and PFC and HPC were dissected on ice. Brain regions were immediately homogenized using pestle tissue grinder in Qiazol before freezing the samples and storage at -80°C. Total RNA was then extracted using the miRNeasy Micro Kit (Qiagen 217084). RNA concentration and purity were verified with a Bioanalyzer 2100 (Agilent Technologies), and all samples had a RIN score above 9.0. RNA libraries were prepared using 200 ng of total RNA following the Illumina Stranded mRNA Ligation Sample Prep Kit protocol. Library concentration and size distribution were assessed using a Bioanalyzer DNA 1000 chip (Agilent Technologies) and Qubit fluorometry (Invitrogen). Libraries were sequenced to obtain 50 M fragment reads per sample, adhering to Illumina’s standard protocol on the NovaSeq™ 6000 SP flow cell. Sequencing was performed as 100 × 2 paired-end reads using the NovaSeq SP sequencing kit and NovaSeq Control Software (version 1.8.0). RNA quality checks and RNA-seq were conducted by the University of Chicago Genomics Core, yielding 30 million paired-end reads via the Illumina NovaSeq 6000, using the oligo dT directional method for library preparation.

Reads were mapped to the mouse reference transcriptome GRCm39, version M27, using STAR (version 2.6.1d). Quality checks were performed with FastQC and MultiQC. Reads were trimmed with trimmomatic package, followed by feature count generation with Rsubread package. Filtering was performed to eliminate non-protein coding genes and genes with low expression across multiple samples (i.e., normalized counts < 5 in more than half of the samples for each sex in for each treatment group). Global differential analyses were carried out for each brain region using DESeq2 version 1.28.1, with fixed effects to account for batch as well as sex and treatment group (Saline+CTRL; MIA+CTRL; Saline+REP; MIA+REP). Analogous differential analyses were also performed for each sex independently, with fixed effects for batch and treatment group. Treatment-specific differentially expressed genes (DEG) between each pair of treatment groups were identified using contrasts and a significance threshold of 10% for the false discovery rate (FDR) K-means clustering was used to cluster normalized expression Z-scores of DEGs to identify modules of distinct transcriptional signatures. Pathway enrichment analysis was performed using clusterProfiler version 4.10.1and Ingenuity Pathway Analysis^®^ software. Upstream regulator analysis was done using Ingenuity Pathway Analysis^®^ software. Data wrangling and graphic generation were conducted using the tidyverse suite of R packages (R version 4.0.0).

### Statistical analysis

All data are presented as means ± standard error of the mean (SEM). Comparisons between groups were done by Student’s *t*-tests. Multiple comparisons were performed by either one-way or repeated measures ANOVA, followed by a Student–Newman– Keuls’s *post hoc* test after assessing for data normality. Data analyses were performed using Prism 8.0 (GraphPad). A statistically significant difference was assumed at p <0.05.

## Results

A recent study demonstrated that a MIA phenotype in offspring depends on the baseline interleukin (IL)-6 reactivity of pregnant female animals to Poly(I:C) (10). In order to strengthen and increase the reproducibility of our MIA model, we built upon the recently published MIA protocol involving the pretesting of the IL-6 immune response of females and the subsequent selection of a susceptible sub-group of females to MIA (10). First, we injected 5 mg/kg of Poly(I:C) intraperitoneally in C57BL/6 (Charles River Laboratory-CRL B6) in non-pregnant female mice and we evaluated their IL-6 blood response 3 hours after the injection followed one week later by the inclusion of the selected sub-group of females into the time-mating pool for the generation of MIA and control offspring (**Fig. 1a**). Based on IL-6 plasma levels, we divided our 7-8 week-old females into low (first quartile), middle (second and third quartile) and high (fourth quartile) responders (**Fig. 1b**). We then injected each category of responders during their pregnancy with two Poly(I:C) doses, 10 or 20 mg/kg, at E9.5 to determine the correct dose to induce a robust MIA phenotype in offspring. E9.5 corresponds to the middle of the first trimester in terms of human developmental biology, at which time studies have reported an increased likelihood of prenatal infection being associated with neurodevelopmental alterations in humans (1). This paradigm has been confirmed to lead to physiological markers of immune response in mothers and microglial alterations in offspring (18). We first evaluated the survival rate of MIA versus saline offspring and found that Poly(I:C) injected pregnant females had a lower survival of litters compared to saline injected females, with the high responder category having the lowest survival of litters (**Fig. 1c, Suppl Table 1**). Based on these results, we decided to exclude the high responder females and focus on the low and middle responder females for sociability evaluation. Here we focused on male offspring to replicate previous studies, including our own (3,4). Social behavior results revealed that only offspring delivered from the middle responder pregnant females injected with 10 mg/kg of Poly(I:C) presented a social behavior deficit among all the middle and low responder females injected with the two Poly(I:C) doses (**Fig. 1d**). Altogether, these data led us to select only the middle responder females and inject a dose of 10mg/kg of Poly(I:C) to induce our MIA model or saline as a control. In addition, knowing that recent papers on MIA reported an important role of IL-17a in the induction of autism-like phenotypes in offspring, associated with the presence of cortical ectopia at E18.5 (11), we checked if it was present in our MIA model. To do that, we assessed C57BL6 from Taconic and C57BL6/J from Jackson laboratory, the two mouse lines used in their paper for cortical ectopia after MIA (11). Both Taconic and JAX B6 pregnant female mice were injected with 10 mg of Poly(I:C) at E12.5 and E9.5 respectively. We could not detect the presence of cortical ectopia during development in either Taconic MIA or Jax MIA offspring at E18.5, but found cortical neuronal layer disorganization in all MIA mice (**Fig. 1e-f**). In regards to IL-17a, we again confirmed its induction in Taconic mice and its absence in Jax mice at 48 hours post injection of Poly(I:C) (**Fig. 1g**).

**Figure 1.**
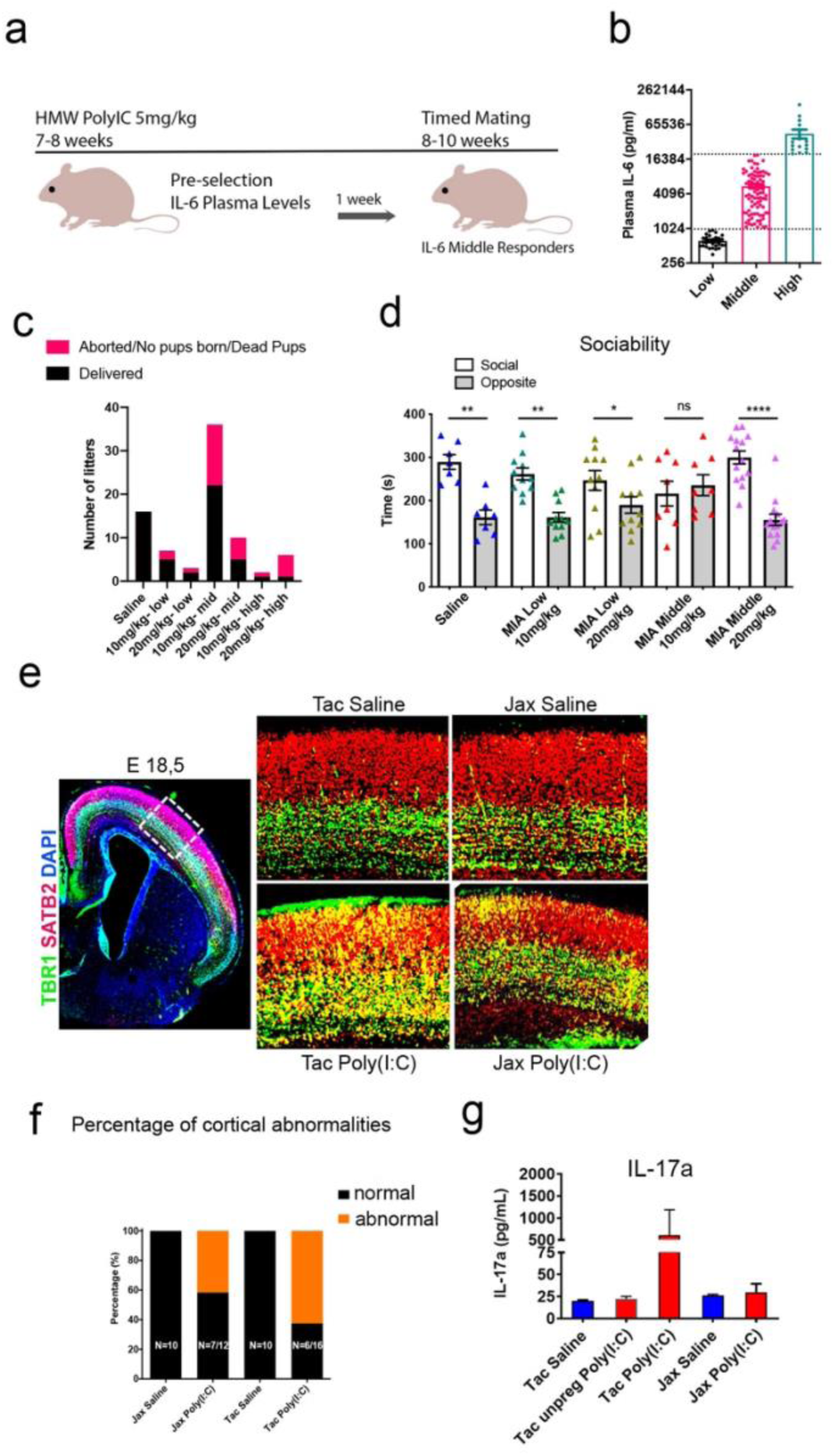
Preselection of female mice for MIA induction based on IL6 response (a) Scheme of the IL-6 pre-selection of females. (b) IL-6 response measured by ELISA 3 hours after HMW Poly(I:C) 5mg/kg injection to establish Low, Middle and High responder females. (c) Graph representing the number of litters with or without delivered pups to establish the success rate of timed mating depending on the Poly(I:C) dose injected in the pregnant females and the category of female responder. (d) Assessment of sociability in male offspring delivered from middle vs low responders using the three-chamber social assay. Social deficits were only present in the offspring from middle responders injected at 10mg/kg at E9.5. (e) Assessment of cortical patch development in the offspring of Taconic (Tac) and Jackson Laboratory (Jax) mice after saline or poly(I:C) treatment by immunofluorescence imaging of upper cortical layer-specific marker, SATB2 (red), deeper cortical layer-specific marker, TBR1 (green), and DAPI (blue) in coronal section of E18.5 frontal cortex. Both Tac Poly(I:C) and Jax Poly(I:C) mouse showed mis-localization of TBR1+ neurons in SATB2+ layer in the somatosensory cortex. Scale bars = 200 um (upper and lower panel). (f) Quantification of the presence of cortical abnormalities in the offspring after saline or Poly(I:C) injection in pregnant Jax and Tax females. (g) Plasma IL-17a levels were examined at 48 hr post-Poly(I:C) ip injection of Tac or Jax female mice at E12.5 for Tac mice and E9.5 for Jax mice. n = (2-5) / group for Tac Saline, Tac Poly(I:C), Jax saline and Jax Poly(I:C). Graph indicates mean±s.e.m.

We then induced our MIA using only middle responder CRL females from the IL6 preselection by injecting them with 10 mg/kg i.p. of Poly(I:C) at embryonic day 9.5 (E9.5), in order to target the initial wave of microglial migration from the yolk sac into the CNS (27), while control females were injected with saline (**Fig. 2a**). We characterized our MIA model in terms of cytokine induction by collecting blood samples from MIA induced pregnant females at 3 and 48 h post injection to analyze plasma IL- 6 (at 3 h) and IL-17 (at 48 h) concentrations by ELISA (**Fig. 2b-c**). IL-6 was significantly increased with middle responders after Poly(I:C) injection compared to the saline injected group (**Fig. 2b**), in agreement with previous literature suggesting IL-6 as one of the primary mediators of maternal inflammation (28). Blood IL-17 levels showed no difference between saline and Poly(I:C) injected female plasma samples (**Fig. 2c**), similar to Jax B6 mice (**Fig. 1g**), ruling out the contribution of IL-17 in our CRL B6 MIA model. The presence of segmented filamentous bacteria was previously reported as necessary to induce maternal IL-17a-dependent-MIA model (29). All of the CRL B6 females used for the timed mating were tested for this bacterial strain and confirmed positive for segmented filamentous bacteria, without inducing IL-17a or cortical ectopia in our MIA model (data not shown). To determine the dynamic of changes in IL-6 level at pre-selection (Poly(I:C) 5 mg/kg, **Fig. 2d**) compared to pregnancy at E9.5 (Poly(I:C) 10 mg/kg, **Fig. 2e**), we performed a paired t-test for Saline and Poly(I:C) (MIA) injected dams. The IL-6 level in saline injected mice at E9.5 was undetectable, as expected (**Fig. 2d**), while the IL-6 level in Poly(I:C) injected mice at E9.5 varied from low to middle range (**Fig. 2e)**, consistent with the recent MIA model developed by the McAllister group (10). Finally, we measured body weight in offspring at P21 and found it significantly lower in male offspring from Poly(I:C) injected dams compared to the saline group, although there was no difference in female offspring between the two groups (**Fig. 2f**).

**Figure 2.**
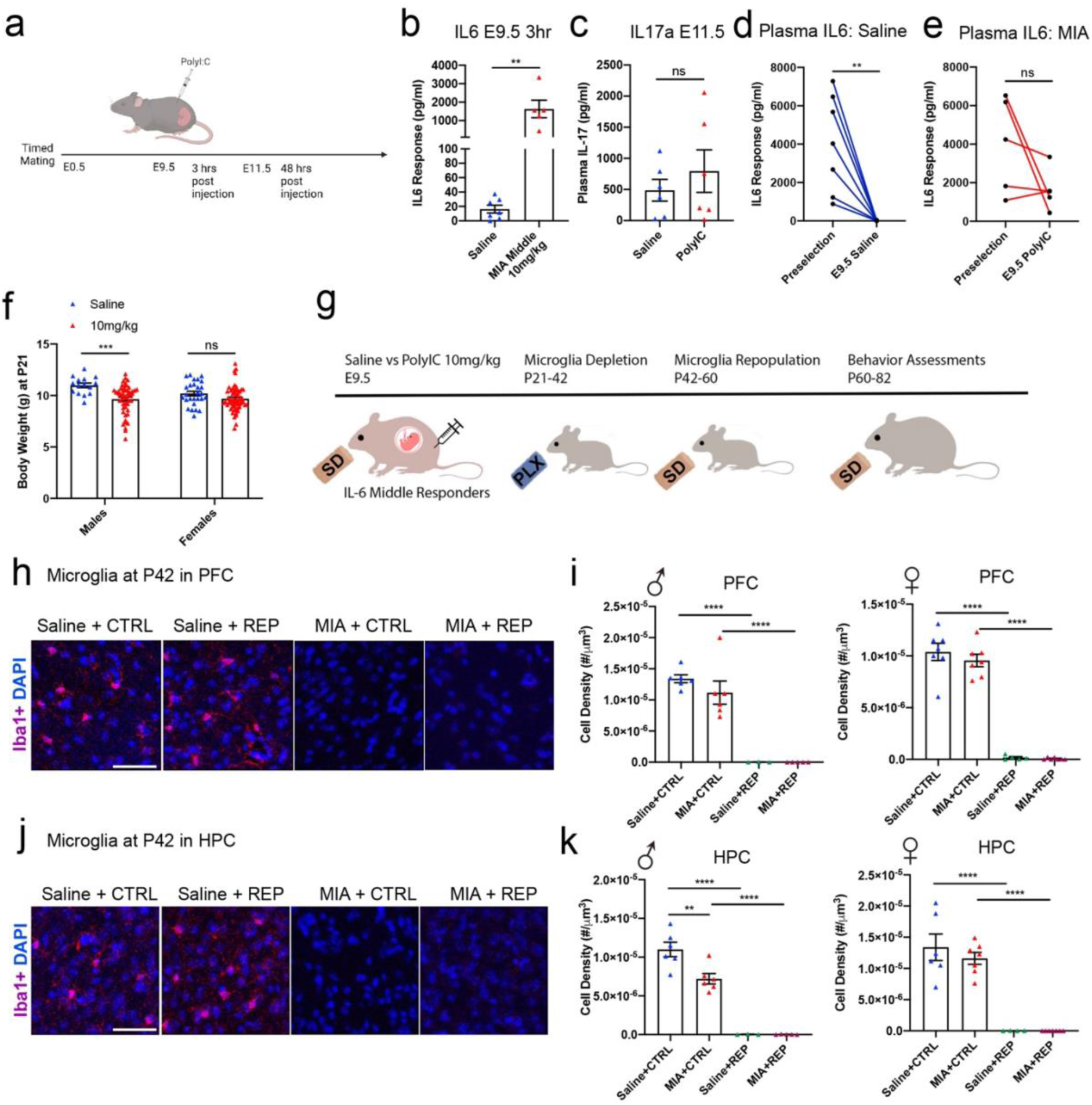
MIA model characterization in middle responder females and evaluation of microglia depletion. **(a)** Scheme indicating the timeline of IL-6 and IL-17 measurements after Poly(I:C) injection at E9.5 in pregnant middle responder females. **(b)** IL-6 response measured by ELISA 3 hours after Poly(I:C) injection at E9.5 in middle responder females. Unpaired t-test. **(c)** IL-17 response measured by ELISA 48 hours after Poly(I:C) injection at E9.5 in middle responder females. Unpaired t-test. **(d)** Graph showing the evolution of plasma IL-6 after Poly(I:C) injection at preselection and saline injection at E9.5. Paired t-test. **(e)** Graph showing the evolution of plasma IL-6 after Poly(I:C) injection at preselection and Poly(I:C) injection at E9.5. Paired t-test. **(f)** Body weight of Saline and MIA offspring at P21. **(g)** Scheme representing the timeline of microglia depletion with CSFR1 inhibitor PLX (*versus* Standard Chow (SD)), renewal and behavior testing after Poly(I:C) injection at E9.5 in middle responder females. **(h-k)** Representative images of microglia staining using antibody against Iba1 at P42 in male and female offspring with or without microglia depletion in PFC **(h)** and HPC **(j)** and the corresponding microglia counting **(I, k)**. Saline or Poly(I:C) injection with control chow (Saline+CTRL ; MIA+CTRL) . Saline or Poly(I:C) injection with PLX5622 treatment (Saline+REP; MIA+REP). Two-way ANOVA, Tukey’s post-hoc, **< 0.01, **** < 0.0001, Graph indicates mean ± s.e.m.

In order to induce microglia depletion followed by spontaneous repopulation, we fed male and female offspring with either control (Saline+CTRL or MIA+CTRL) or PLX5622 to induce microglial depletion between postnatal days 21-42, and then replaced PLX5622 chow with normal chow, leading to microglia replenishment starting at P42 (Saline+REP or MIA+REP, **Fig. 2g**) as previously published (4,19). We evaluated microglia presence in the PFC and HPC of offspring after MIA and PLX5622 treatment at P42 by immunofluorescence using antibody directed against Ionized calcium-binding adapter molecule 1 (Iba1). We found a significant reduction in microglia number by MIA only in male HPC and confirmed the depletion of Iba1^+^ microglia at P42 in both regions in male and female offspring (**Fig. 2h-k)**. We can therefore confirm that PLX5622 treatment significantly reduced microglia number in both regions and that there was no difference in the treatment effect by sexes.

We next evaluated behavioral phenotypes using a series of tests to assess social, sensorimotor, learning, memory and emotional dimensions in offspring. We tested them in the following order: social chamber, open field, Y-maze forced alternation, novel object recognition, pre-pulse inhibition (PPI) and fear conditioning test between P60-82 (**Fig. 3a**). There was no change in the time spent in the center of the open field, or the distance and velocity among the 4 treatment groups including males and females, suggesting MIA or PLX5622 has no effect on anxiety-like behavior, locomotion or exploratory behavior at adulthood (**Suppl Fig. 1a-c)**. In order to assess the canonical neurodevelopmental disorder-related social dimension in adult offspring, we performed the three-chamber test at P60-63 to measure the effects of MIA and PLX on sociability and social novelty in both males and females by measuring the total time spent in the social versus opposite empty chamber and in the new mouse versus opposite familiar mouse-containing chamber, respectively. Sociability depends on PFC and subcortical areas for the reward aspect of social interaction (30), while social novelty recognition depends more on hippocampus and central and medial amygdala (31–33). We found that MIA induced deficits for both sociability and social novelty with MIA+CTRL male offspring compared to Saline+CTRL male offspring mice (**Fig. 3b-c**). PLX5622 treatment of male mice significantly normalized sociability at adulthood **(Fig. 3b, left)**, reproducing our previous results published in *Mol Psychiatry* (4), while there was no effect on social novelty **(Fig. 3c, left).** We further observed a deficit in sociability in MIA female mice (**Fig. 3b, right**), which was not normalized by PLX5622 treatment, suggesting a sex-specific therapeutic effect of microglial depletion on sociability deficit. Finally, female mice showed normal social novelty (**Fig. 3c, right**), indicating a sex specific MIA phenotype in regards to social novelty recognition and suggesting an alteration in medial amygdala in MIA males not corrected by PLX (**Suppl Fig. 1e**).

**Figure 3.**
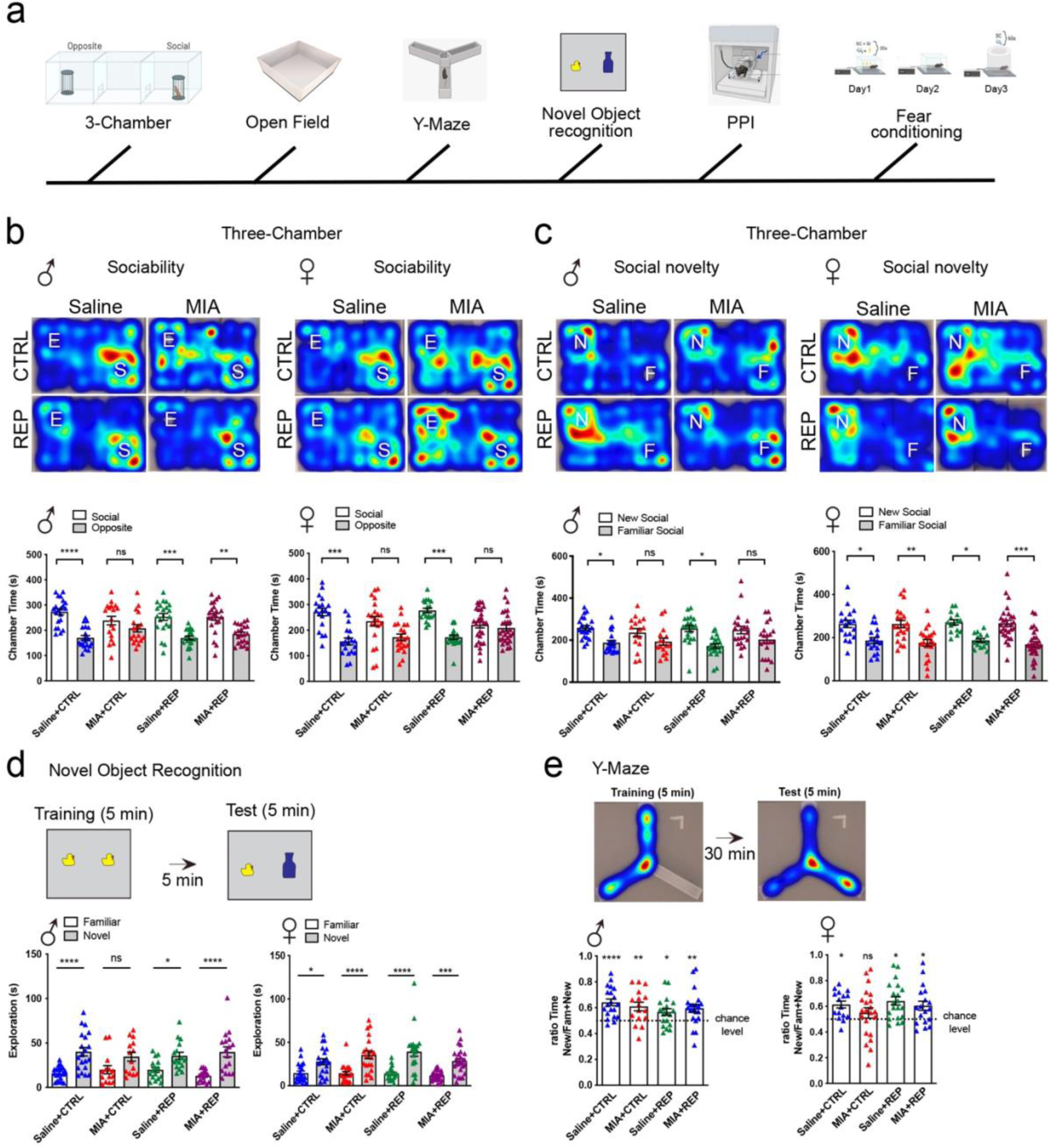
Assessment of social and short-term memory abilities in MIA offspring. **(a)** Schematic timeline of the behavior battery conducted. **(b-c)** Three chamber social assay representative tracking images of sociability **(b)** and social novelty **(c)** evaluation. Sociability deficit induced by MIA was corrected in MIA+REP group by PLX5622 treatment in both male (left graph) and female (right graph) **(b)**. Social novelty was affected in male (left graph) without correction in MIA+REP group by PLX5622 treatment. No difference in social novelty in female (right graph) **(c)**. **(d)** Scheme of the novel object recognition paradigm showing the presence of a novel object during the test session (left). Time exploring the familiar and novel objects showing no differences among the groups in males (center graph) and females (right graph). **(e)** Scheme of the Y-maze test showing the novel arm of the maze to be explored during the test session. Ratio of the time exploring the novel arm over the novel and familiar arms in male showing no differences among the groups (center graph), and in female showing a deficit in MIA animals corrected in MIA+REP group by PLX5622 treatment (right graph). Two-way ANOVA, Tukey’s post-hoc, *<0.05, **< 0.01, ***< 0.001, **** < 0.0001, Graph indicates mean±s.e.m.

Next, we aimed to determine the effects of MIA and PLX on learning and memory abilities at adulthood. We first examined if MIA interferes with working memory and used the object recognition test, which is mainly regulated by the perirhinal cortex and hippocampus (34). Male mice from the MIA+CTRL group showed a deficit in novel object recognition compared to the Saline+CTRL group and a recovery after PLX5622 treatment. On the contrary, no effect of MIA nor PLX5622 was found in the novel object recognition in female mice (**Fig. 3d**). Mice were next examined for their spatial working memory by a forced alternation Y-maze test at P70. MIA+CTRL male mice showed normal working memory and overall, all male mice showed correct spatial working memory in this test. On the contrary, MIA+CTRL female mice showed deficient working memory, which was normalized by PLX5622 treatment (**Fig. 3e**). Taking the two tests assessing working memory abilities into account and looking only at saline groups, no effect of PLX treatment by itself was found. We then examined the sensorimotor gating abilities of the MIA and control offspring. A pre-pulse inhibition (PPI) test is widely used to measure deficits in information-processing abilities or sensorimotor gating across species, including humans and rodents (35). Mice were tested for PPI at P77, and we reported no deficit in PPI in any group in both male and female offspring **(Suppl Fig. 1d**), suggesting normal sensory gating abilities in MIA offspring and no effect of PLX treatment by itself (**Suppl Fig. 1e**).

Finally, mice were tested for associative learning memory at P80-82 using the fear conditioning paradigm. Males and females in all four groups showed normal learning abilities during the acquisition phase on day 1 (**Fig. 4a**). On day 2, we found significantly reduced contextual fear memory in MIA+CTRL male and female offspring, suggesting that MIA induced hippocampal-dependent memory impairment in both sexes. Strikingly, reduction in contextual memory was corrected by PLX5622 treatment in MIA+REP male offspring. MIA+REP female was similar to Saline+REP female offspring with REP female offspring showing overall lower freezing compared to CTRL offspring. These data indicate the potential microglial involvement in MIA-mediated hippocampal impairment (**Fig. 4b**). Cued memory, whose acquisition and consolidation is known to be regulated by the amygdala (36), was normal among the four groups on day 3 for both sexes and unaffected by PLX5622 treatment (**Fig. 4c**), suggesting that amygdala neuronal network, including the basolateral amygdala, involved in cued- associative memory (37) was unaffected by MIA or PLX treatment (**Suppl Fig. 1e**).

**Figure 4.**
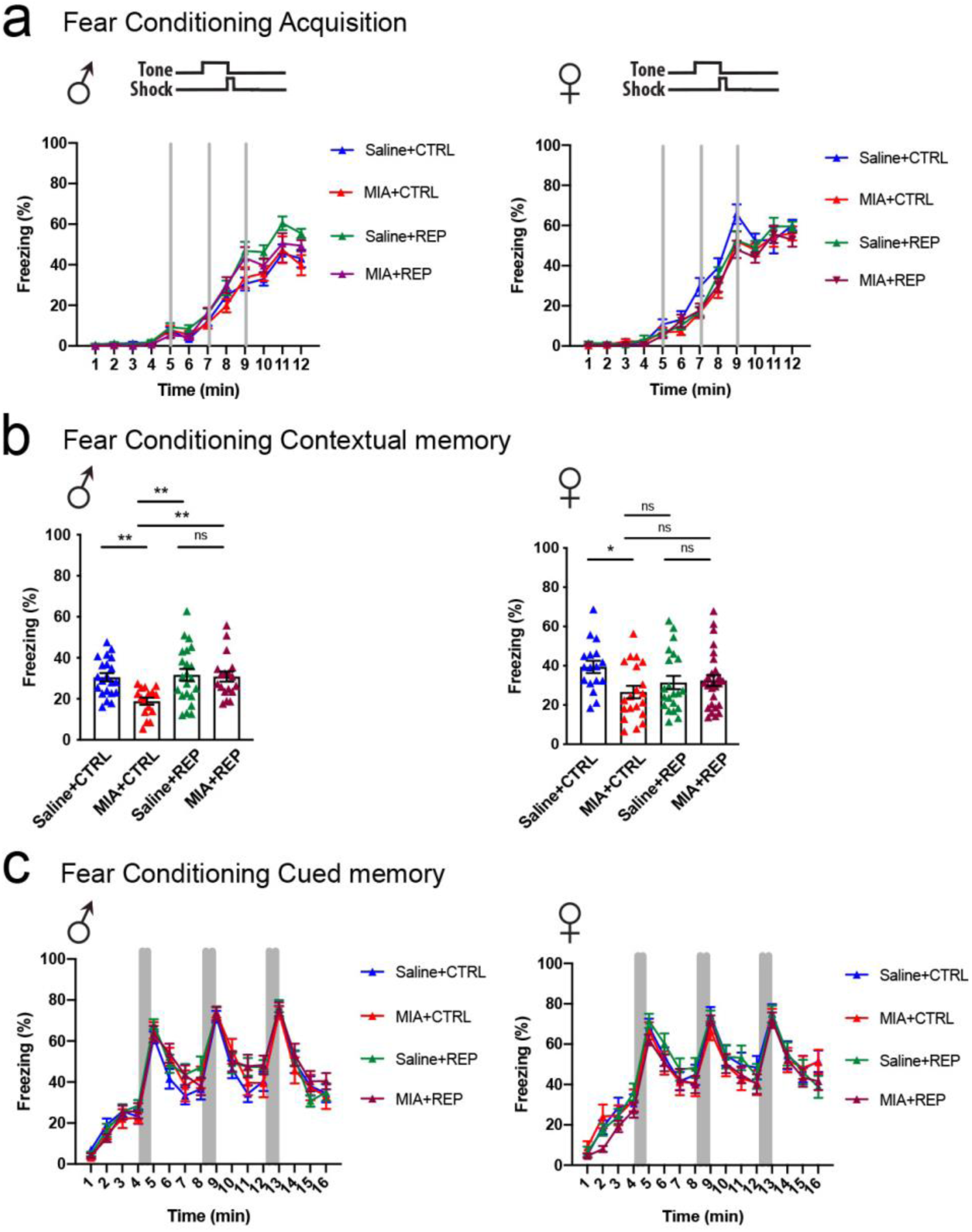
Assessment of long-term memory abilities in MIA offspring. **(a)** Time-course of freezing observed in the fear conditioning paradigm during training on day 1 showing no differences among the groups in males (left graph) and females (right graph). **(b)** Average freezing time observed during contextual memory testing on day 2 in males (left graph) and females (right graph). **(c)** Time-course of freezing observed during cued memory testing on day 3 in males (left graph) and females (right graph). Graph indicates mean±s.e.m.

To study the sex differences related to PLX treatment in individuals with MIA, with an aim to better understand the mechanisms linked to the detrimental effects of MIA and beneficial effects of our microglial renewal approach, we used RNA-seq analysis of mouse brain structures at adulthood. We performed transcriptomic analysis of the PFC and HPC homogenates of Saline+CTRL; MIA+CTRL; Saline+REP and MIA+REP male and female offspring at P90. We focused on PFC and HPC as these two structures are primarily involved in sociability (social dimension) (38) and spatial memories (learning and memory dimension) (39), respectively, and our behavioral data indicate a detrimental effect of MIA treatment in both paradigms, which were partially corrected by PLX in a sex-specific manner. We first analyzed the transcriptomic profile of HPC to gain knowledge on the neurobiology at play in regards to learning and memory abilities. *K*-means clustering (*k*=3) of 697 differentially expressed genes (DEGs) from combined male and female RNA-seq data revealed distinct transcriptional signatures enriched by PLX (violet module, 219 genes), downregulated by MIA and corrected by PLX (red module, 253 genes), upregulated by MIA and corrected by PLX (yellow module, 225 genes). (**Fig. 5a; Suppl Table 2**). Gene ontology (GO) enrichment analysis of the biological processes of each module indicates that the PLX effect, represented in the violet module, is mainly related to nuclear changes, potentially related to the rejuvenated status of the myeloid cells in the brain after PLX treatment (**Fig. 5b; Suppl Table 2**). More importantly, the enrichment analysis of the red module, representing the genes downregulated by MIA and corrected by PLX, highlights terms related to metabolic changes (**Fig. 5b; Suppl Table 2**) and the yellow module, representing the genes enriched by MIA and corrected by PLX, highlights learning and memory terms, both potentially explaining the beneficial effects of PLX on spatial memories in both sexes (**Fig. 5b; Suppl Table 2**). To distinguish the mechanisms involved, we looked at the specific DEGs per sex, affected by MIA in comparison to all the other groups (**Fig. 5c-d)**. We found more DEGs in females compared to males with an overall higher number of downregulated genes (**Fig. 5d**). We found that most of these DEGs are sex-specific, with only 8 DEGs in common between males and females in the MIA+CTRL vs all remaining groups (ALL) comparison. In females and males, respectively, we also found that 18 and 9 DEGs were affected in both the MIA+CTRL vs ALL and MIA+CTRL vs MIA+REP comparisons (**Fig. 5c; Suppl Table 2**).

**Figure 5.**
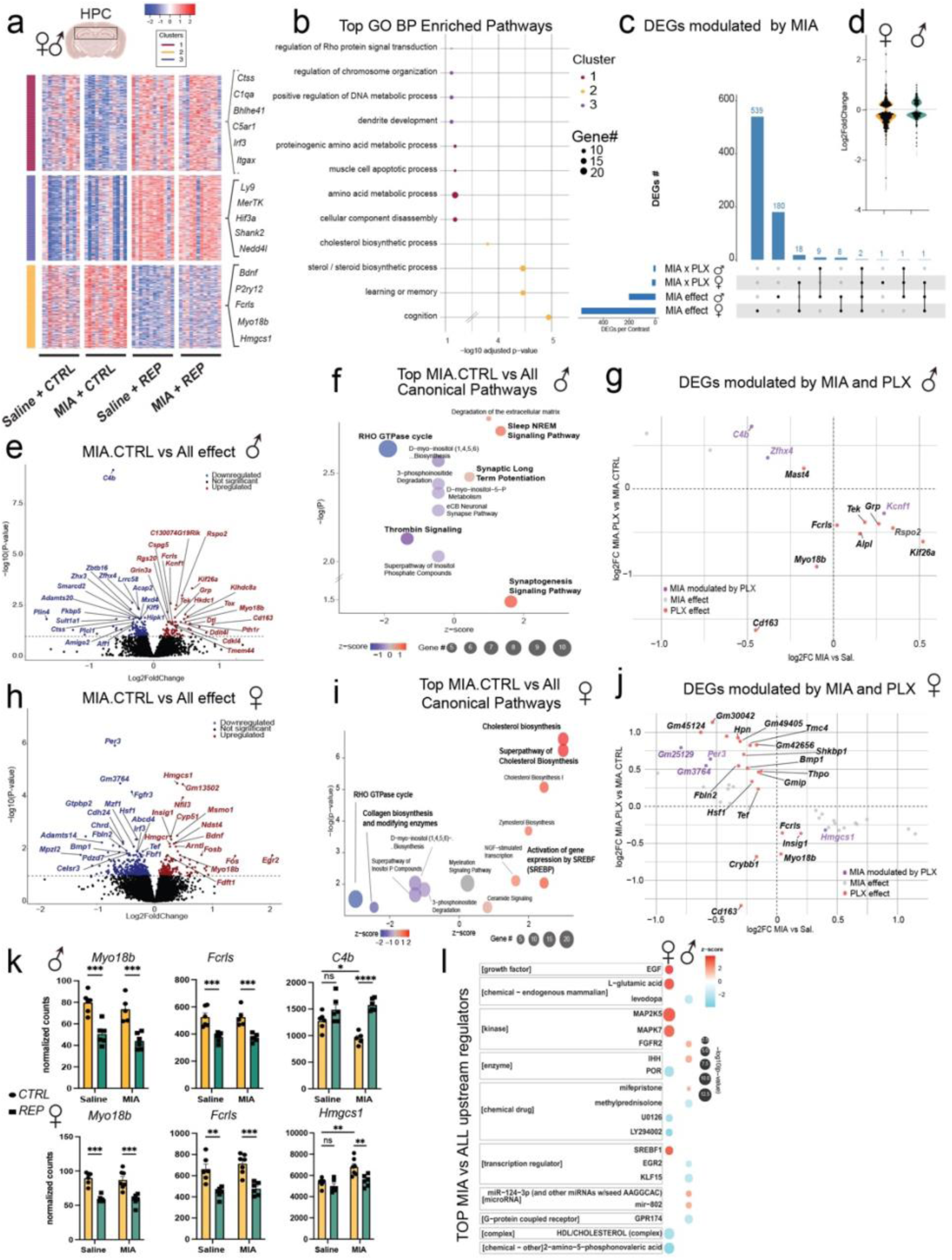
MIA alters hippocampal gene pathways in a sex-specific manner and is modulated by PLX. **(a)** Heatmap of differentially expressed hippocampal genes (DEGs) from mice that received Saline (Saline+CTRL) or Poly(I:C) (MIA+CTRL) injection, +/- PLX5622 treatment (Saline+REP; MIA+REP) (*n* = 6 mice per group, DESeq2, FDR<0.1). **(b)** Top-enriched Gene Ontology Biological Processes pathways for each cluster from (A). **(c)** Distribution of MIA DEGs per sex, +/- PLX5622 treatment. MIA x PLX = MIA+CTRL VS MIA+REP contrast. MIA effect = MIA+CTRL VS ALL contrast. **(d)** Log2 fold change of MIA affected DEGs in females vs. males. **(e)** Volcano plot of MIA DEGs in males (*n* = 6 per group). **(f)** Top-affected canonical pathways in males from IPA are represented. **(g)** Scatter plot representing MIA DEGs from male mice, that are modulated by PLX5622 treatment (FDR<0.1). **(h)** Volcano plot of MIA DEGs in females (*n* = 6 per group). **(i)** Top-affected canonical pathways in females from IPA are represented. **(j)** Scatter plot representing MIA DEGs from female mice, that are modulated by PLX5622 treatment (FDR<0.1). **(K)** Representative bar plots of normalized counts for commonly modulated DEGs in both male and female mice after PLX5622 treatment and specific DEG affected by MIA and corrected by PLX in male and female mice. **(l)** Top MIA Upstream regulators from IPA in both male and female MIA mice. Graph indicates mean±s.e.m.

We then focused on DEGs specifically affected in the comparison of MIA+CTRL vs ALL split by sex. Male DEGs affected by MIA in comparison to all of the other groups included 79 upregulated and 124 downregulated genes (**Fig. 5e; Suppl Table 2**). Among these, Ingenuity Pathway Analysis showed an upregulation of canonical pathways including Synaptic long-term potentiation and Synaptogenesis signaling and downregulation of metabolic and RHO GTPase cycle IPA-enriched pathways (**Fig. 5f; Suppl Table 2**). In particular, a comparison of MIA vs. Sal log-fold changes against MIA+REP vs. MIA+CTRL log-fold changes highlighted DEGs specifically impacted by PLX, including *fcrls* and *myo18b*, as well as genes affected by MIA and corrected by PLX in male HPC, including *C4b* (**Fig. 5g, k; Suppl Table 2**). Female DEGs affected by MIA in comparison to all other groups included 186 upregulated and 382 downregulated genes (**Fig. 5h; Suppl Table 2**), corresponding to upregulation of lipid metabolism pathways and downregulation of metabolic and RHO GTPase cycle pathways, consistent with the male dataset (**Fig. 5i; Suppl Table 2**). A comparison of MIA+CTRL log-fold changes against those of MIA+REP vs. MIA+CTRL highlighted DEGs specific to the PLX effect, including *fcrls* and *myo18b*, and genes affected by MIA and corrected by PLX in female HPC, including *Hmgcs1* (**Fig. 5j, k; Suppl Table 2**). Interestingly, downregulated pathways are similar between males and females, whereas upregulated pathways are different by sex. Finally, upstream regulator analysis identified SREBP1 as a transcription factor upregulated in MIA females and EGR2 and KLF15 as downregulated transcription factors in MIA males (**Fig. 5l; Suppl Table 2**).

In regards to PFC transcriptomic analysis, *K*-means clustering (*k*=3) of 902 differentially expressed genes (DEGs) from combined male and female RNA-seq data revealed distinct transcriptional signatures enriched by PLX (violet module, 288 genes), downregulated by PLX (red module, 394 genes), upregulated by MIA and reversed by PLX (yellow module, 220 genes). (**Suppl Fig. 2a; Suppl Table 2**). GO enrichment analysis of the biological processes of each module indicates that the PLX effect, represented by the violet and red modules, is mainly related to metabolic changes, resembling HPC transcriptomic results. The yellow module, representing genes affected by MIA and reversed by PLX, revealed enrichment of molecules related to cellular differentiation (**Suppl Fig. 2b; Suppl Table 2**). Most of the DEGs were found to be unique per sex, with only 4 DEGs in common for both sexes in the MIA+CTRL vs ALL contrast. We also found that in females and males, respectively, 80 and 5 DEGs were affected in both the MIA+CTRL vs ALL and MIA+CTRL vs MIA+REP comparisons (**Suppl Fig. 2c; Suppl Table 2**). Of note, the number of DEGs significantly affected per sex was higher in females (824 genes) compared to males (37 genes) (**Suppl Fig. 2d; Suppl Table 2)**. When focusing on DEGs specifically affected in the MIA+CTRL vs ALL comparison split by sex, we found 16 genes upregulated and 21 downregulated in males for MIA in comparison to all other groups (**Suppl Fig. 2e**), corresponding to two IPA-enriched pathways including downregulation of phagosome (**Suppl Fig. 2f; Suppl Table 2**). Comparisons of MIA+CTRL log-fold changes against those of MIA+REP vs. MIA+CTRL revealed DEGs affected either by MIA or PLX in males, including *fcrls, myo18b* and *Napepld* (**Suppl Fig. 2g, k; Suppl Table 2**). Female DEGs affected by MIA in comparison to all other groups represent 370 upregulated and 457 downregulated genes (**Suppl Fig. 2h; Suppl Table 2**), corresponding to upregulation of SNARE signaling pathway and downregulation of extracellular matrix and RHO GTPase cycle pathways (**Suppl Fig.**

**2i; Suppl Table 2**). A comparison of MIA+CTRL log-fold changes against those of MIA+REP vs. MIA+CTRL highlighted DEGs specific to PLX effect, including as in HPC *fcrls* and *myo18b* and only three genes affected by MIA and corrected by PLX in female PFC, including *C4a* (**Suppl Fig. 2j, k; Suppl Table 2**). Finally, upstream regulator analysis identified mainly RICTOR, a potential master regulator of mTOR pathway, as a downregulated transcription factor in MIA females, indicated upregulation of pathways related to epilepsy (bicuculline and kainic acid regulators) and only weak identifications in MIA males (**Suppl Fig. 2l; Suppl Table 2**).

## Discussion

We demonstrated sex-specific MIA-mediated abnormal behaviors, including cognitive impairment and social deficits in MIA offspring (**Suppl Fig. 1e**). In addition, we showed that CSF1R inhibitor corrected MIA-induced behavior deficits at young adult phase in a sex-specific manner. With pre-selection of CRL B6 female mice based on baseline immunoreactivity to low dose Poly(I:C) stimulation (10), we identified the middle- responders as the optimal responders for MIA and 10mg/kg of our lot of Poly(I:C) as the optimal dose for MIA induction at E9.5 to induce abnormal behaviors in the offspring with an average of 39% litter loss. The variability of baseline immunoreactivity and immune response to MIA induction is a challenge for studying MIA, which makes comparing the outcomes between studies very difficult (10). We stress the importance of taking into account baseline immunoreactivity of the future mothers and the importance of using Poly(I:C) with the same catalog number and lot number to minimize variability in order to obtain robust and reproducible abnormal behaviors in both sexes (10).

Sex-specific effects of MIA on social behavior and cognition have been previously reported. *Xuan et al* showed increased repetitive behavior only in male offspring mice from dams injected with Poly(I:C) at E12.5, while no sex differences were observed for social deficits induced by MIA (40). Another study also demonstrated a male-specific MIA effect by showing reduced sociability only in male offspring from Poly(I:C) injected dams at E12.5 (41). Recently, Gogos *et al* found spatial working memory deficits only in males and reduced pre-pulse inhibition and increased acute methamphetamine-induced locomotor hyperactivity in both sexes of rat offspring from dams injected with Poly (I:C) at E15 (42). These previous studies reveal MIA-induced deficits in only males in certain behavior assays.

Previous studies were based on a single Poly(I:C) injection without pre-selection while Estes *et al* also reported a male-specific MIA effect on repetitive behavior after the pre-selection and MIA induction with Poly(I:C) injection (30mg/kg) in mice at E12.5. Here we found MIA-induced social deficits and spatial memory impairments associated with dysfunction in PFC and HPC in both sexes, while there was a sex-specific deficit in social novelty recognition in male offspring, which relies notably on the central and medial amygdala (31,33,43). No effects on anxiety or cued-associative memory, which relies notably on the basolateral amygdala (36,37,44), were observed, suggesting that MIA may affect specific amygdala nuclei. Sensorimotor gating abilities were not affected by MIA, suggesting proper information processing and no attentional deficit in the sensorimotor system of male and female offspring. The time frame or dose of the Poly(I:C) injection to induce MIA may affect specific brain regions or recruit the involvement of different cell types in the pathological process underlying MIA (10). Importantly, the dose of Poly(I:C) also depends on the lot, as higher doses of Poly(I:C) at 20mg/kg led to no MIA phenotype in offspring in this study (**Fig. 1d**), whereas 20mg/kg Poly(I:C) from a different lot at E9.5 without pre-selection induced social deficit in male offspring in our previous work (4). Determination of the efficient dose according to the Poly(I:C) lot is thus crucial, supporting the observation of many differences among different lots (45). Further studies are still required to understand the precise mechanisms contributing to the variable outcomes for abnormal behaviors in MIA offspring.

We have previously reported reversed social deficit in male offspring by administering CSF1R inhibitor followed by microglia replenishment (4). We observed in this study similar therapeutic effects of CSF1R inhibition and microglia repopulation in MIA offspring delivered from pre-selected middle inflammatory responders females. CSF1R inhibition followed by microglia repopulation treatment rescued sociability deficit in males, spatial working and associative memory impairments in both males and females, suggesting that microglia play a critical role in mediating MIA-induced abnormal behaviors in social and spatial memory dimensions. There was no sex difference in the pharmacological response to CSF1R inhibitor as the drug depleted microglia nearly 100% in both males and females. Lack of therapeutic effect of CSF1R on sociability deficit in females indicates a sex-specific mechanism of MIA-mediated abnormal behavior, which may be rescued by targeting a different molecule or cell type.

The lack of correction of social novelty recognition abnormality found in males indicates that not all MIA-induced behavioral deficits can be corrected by a microglial renewal strategy. Based on our results, the therapeutic effect of CSF1R during young adulthood could be limited to sociability and spatial memory dimensions (**Suppl Fig 1.e**). Further work using this approach is needed to clearly delineate the beneficial effect of microglial renewal on MIA-induced phenotype using different time windows of CSF1R treatment, longer time after CSF1R treatment, and different MIA models.

Taking all of our results into account, it is important to note that transient PLX5622 treatment does not alter any of the behavioral dimensions assessed in both sexes, indicating no adverse effect after withdrawal of this drug per se. This is aligned to previous data showing that chronic microglia elimination using PLX5622 drug increased neuronal activity and that after microglia repopulation these effects were gone (46). Given that we reported the presence of a distinct transcriptomic signature induced by CSF1R treatment in both PFC and HPC, complementary measures at later time point would be important to determine potential long-term adverse effects of such a microglial renewal approach.

Our brain structure bulk transcriptomic results highlighted specific MIA effects per sex, with a more pronounced effect in females compared to males in both PFC and HPC. The identification of pathways related to cognition, learning and memory abilities in HPC affected by MIA and reversed by PLX is in agreement with the spatial memory impairments we found and its correction after PLX treatment. Male offspring specific pathway analysis indicated the possible involvement of synaptogenesis related mechanisms, which is in agreement with the literature reporting a key role of microglia in synaptic refinement (13) and with previous work showing that PLX treatment can restore microglial-related synaptic contact abnormalities in MIA offspring (4). In female offspring specifically, the identification of lipid-related metabolic pathways and notably the upregulation of the cholesterol biosynthesis pathway is highly interesting as hypocholesterolemia has been reported to be linked to autism and genetic syndromes of neurodevelopmental disorders in human (47). Further research would be necessary to evaluate cholesterol levels in MIA offspring and determine if the upregulation of the cholesterol biosynthesis pathway could be due to a possibly more pronounced hypocholesterolemia status in MIA female offspring. In regards to PFC, the small number of genes affected in males limits our ability to identify a relevant pathway at adulthood involved in the MIA and PLX effects, especially since no DEGs affected by MIA and corrected by PLX were identified. Female PFC results indicate alteration of ECM and the mTOR pathways (48), and potential upregulation of pathways related to epilepsy (49) in MIA without a correction by PLX. A recent hypothesis linked ECM component internalization to cellular metabolism, though further research is needed to define the molecular mechanisms controlling ECM trafficking and fully elucidate how this pathway impinges on cellular metabolism (50). However, our approach does rely on the bulk transcriptomics, thus limiting our ability to determine the effects of MIA and PLX specific to certain cell types. Future experiments using single cell sequencing or proteomic approaches would greatly complement our insight into MIA effects on behavioral dimensions and their improvement with CSF1R treatment.

Overall, we reported no adverse effect on learning and memory abilities induced by PLX5622 treatment and a beneficial effect of PLX5622 treatment on sociability, working and associative memory abilities at adulthood in both sexes. We thus demonstrated that a microglial renewal approach at early adulthood is a promising strategy to correct MIA-induced social and spatial memory ability deficits at adulthood in both sexes.

## Author Contributions

Conceptualization, JC.D., S.I. and T.I.; methodology, JC.D., H.Y.; formal analysis, JC.D., H.Y., P.M., C.M, A.R. ; writing original draft preparation, JC.D., H.Y., C.M., S.I., T.I.; writing—review and editing, S.I., T.I., A.R.; visualization, JC.D., H.Y., P.M., C.M.; supervision, S.I. and T.I.; funding acquisition, T.I, S.I., and JC.D. All authors have read and agreed to the published version of the manuscript.

## Institutional Review Board Statement

All animal procedures followed the guidelines of the National Institutes of Health Guide for the Care and Use of Laboratory Animals and were approved by the Boston University Institutional Animal Care and Use Committee.

## Data Availability Statement

The RNA-seq dataset that support the findings of this study will be deposited in the Gene Expression Omnibus (GEO).

## Acknowledgments

We would like to thank Dr. Clarence Schutt at the Nancy Lurie Marks Family Foundation, Robert Landreth at the Landreth Family Fund and Dr. Susan Leeman at Boston University School of Medicine for supporting this study. We also would like to thank Dr K. McAllister for precious exchanges in regard to the IL-6 response for MIA preselection model and Plexxikon, Inc. for providing PLX5622.

---

**Supplementary Figure 1.**
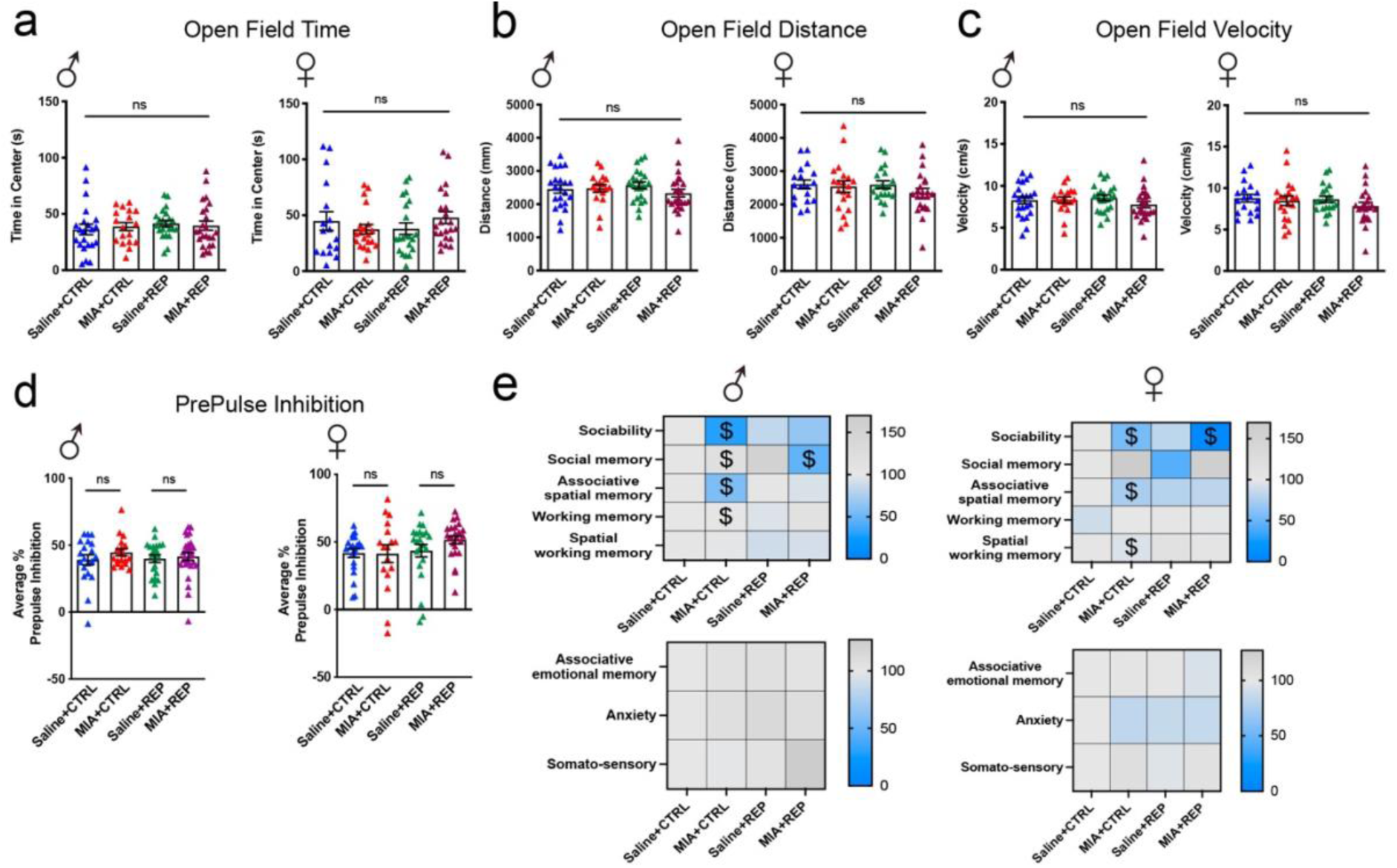
Assessment of anxiety, locomotor activity and sensorimotor gating in MIA offspring. **(a-c)** Open field test evaluating anxiety level **(a)** and locomotor activity **(b-c)** in males (left graphs) and females (right graphs) shows no difference among the groups. **(d)** Average percentage of prepulse inhibition of the startling response showing no differences among the groups in males (left graph) and females (right graph). **(e)** Heatmaps representing a summary of the behaviors affected by MIA and corrected by PLX in males (Top-Left) and females (Top-Right) and heatmaps of the non-affected behaviors at the bottom. $ indicates significative difference compared to Saline+CTRL group. Graph indicates mean±s.e.m.

**Supplementary Figure 2.**
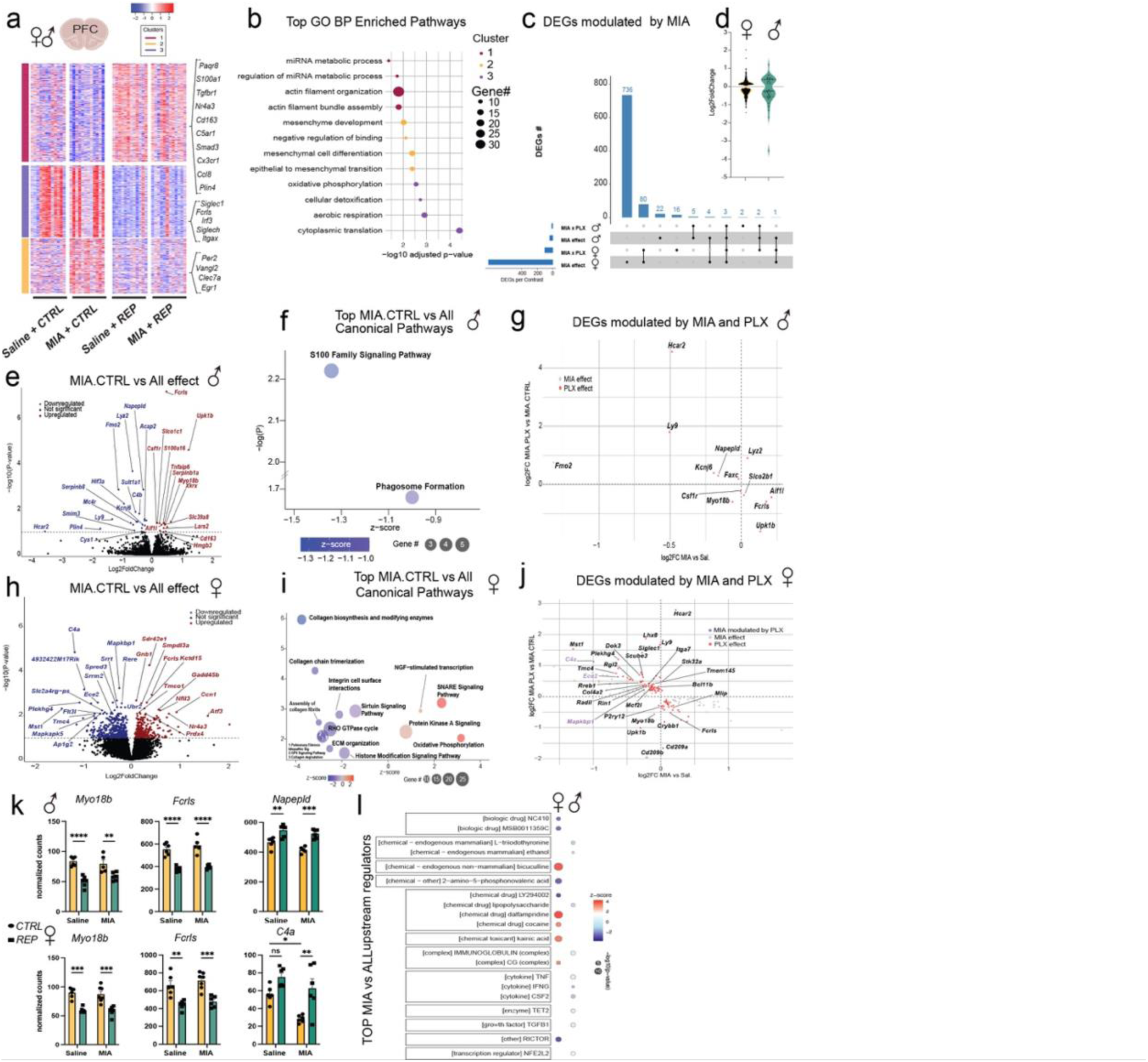
MIA alters prefrontal cortex gene pathways in a sex-specific manner and is modulated by PLX. **(a)** Heatmap of differentially expressed prefrontal cortex genes (DEGs) from mice that received Saline (Saline + CTRL) or Poly(I:C) (MIA + CTRL) injection, +/- PLX5622 treatment (Saline + REP; MIA + REP) (*n* = 6 mice per group, DESeq2, FDR<0.1). **(b)** Top-enriched Gene Ontology Biological Processes pathways for each cluster from (A). **(c)** Distribution of MIA DEGs per sex, +/- PLX5622 treatment. MIA x PLX = MIA+CTRL VS MIA+REP contrast. MIA effect = MIA+CTRL VS ALL contrast. **(d)** Log2 fold change of MIA affected DEGs in females vs. males. **(e)** Volcano plot of MIA DEGs in males (*n* = 6 per group). **(f)** Top-affected canonical pathways in males from IPA are represented. **(g)** Scatter plot representing MIA DEGs from male mice, that are modulated by PLX5622 treatment (FDR<0.1). **(h)** Volcano plot of MIA DEGs in females (*n* = 6 per group). **(i)** Top-affected canonical pathways in females from IPA are represented. **(j)** Scatter plot representing MIA DEGs from female mice, that are modulated by PLX5622 treatment (FDR<0.1). **(k)** Representative bar plots of normalized counts for commonly modulated DEGs in both male and female mice after PLX5622 treatment. **(l)** Top MIA Upstream regulators from IPA in both male and female MIA mice. Graph indicates mean±s.e.m.

**Supplementary Table 1.** Survival rate of MIA versus saline offspring based on Poly(I:C) dose of injection. Survival rate determined based on number of successful deliveries divided by number of pregnant dams injected with Poly(I:C).

|  | MIA 10mg/kg |  |  |  | MIA 20mg/kg |  |  |  | Saline |  |  |  |
| --- | --- | --- | --- | --- | --- | --- | --- | --- | --- | --- | --- | --- |
|  | Low | Middle | High | MIA Total | Low | Middle | High | MIA Total | Low | Middle | High | Saline Total |
| Number of injection | 7 | 36 | 2 | 45 | 3 | 10 | 6 | 19 | 4 | 8 | 4 | 16 |
| Number of delivery | 5 | 22 | 1 | 28 | 2 | 5 | 1 | 8 | 4 | 8 | 4 | 16 |
| Survival Rate | 71% | 61% | 50% | 62% | 67% | 50% | 17% | 42% | 100% | 100% | 100% | 100% |

